# Controlling for Human Population Stratification in Rare Variant Association Studies

**DOI:** 10.1101/2020.02.28.969477

**Authors:** Matthieu Bouaziz, Jimmy Mullaert, Benedetta Bigio, Yoann Seeleuthner, Jean-Laurent Casanova, Alexandre Alcais, Laurent Abel, Aurélie Cobat

## Abstract

Population stratification is a strong confounding factor in human genetic association studies. In analyses of rare variants, the main correction strategies based on principal components (PC) and linear mixed models (LMM), may yield conflicting conclusions, due to both the specific type of structure induced by rare variants and the particular statistical features of association tests. Studies evaluating these approaches generally focused on specific situations with limited types of simulated structure and large sample sizes. We investigated the properties of several correction methods in the context of a large simulation study using real exome data, and several within- and between- continent stratification scenarios. We also considered different sample sizes, with situations including as few as 50 cases, to account for the analysis of rare disorders. In this context, we focused on a genetic model with a phenotype driven by rare deleterious variants well suited for a burden test. For analyses of large samples, we found that accounting for stratification was more difficult with a continental structure than with a worldwide structure. LMM failed to maintain a correct type I error in many scenarios, whereas PCs based on common variants failed only in the presence of extreme continental stratification. When a sample of 50 cases was considered, an inflation of type I errors was observed with PC for small numbers of controls (≤100), and with LMM for large numbers of controls (≥1000). We also tested a promising novel adapted local permutation method (LocPerm), which maintained a correct type I error in all situations. All approaches capable of correcting for stratification properly had similar powers for detecting actual associations pointing out that the key issue is to properly control type I errors. Finally, we found that adding a large panel of external controls (*e.g*. extracted from publicly available databases) was an efficient way to increase the power of analyses including small numbers of cases, provided an appropriate stratification correction was used.

**Author Summary:** Genetic association studies focusing on rare variants using next generation sequencing (NGS) data have become a common strategy to overcome the shortcomings of classical genome-wide association studies for the analysis of rare and common diseases. The issue of population stratification remains however a substantial question that has not been fully resolved when analyzing NGS data. In this work, we propose a comprehensive evaluation of the main strategies to account for stratification, that are principal components and linear mixed model, along with a novel approach based on local permutations (LocPerm). We compared these correction methods in many different settings, considering several types of population structures, sample sizes or types of variants. Our results highlighted important limitations of some classical methods as those using principal components (in particular in small samples) and linear mixed models (in several situations). In contrast, LocPerm maintained a correct type I error in all situations. Also, we showed that adding a large panel of external controls, *e.g* coming from publicly available databases, is an efficient strategy to increase the power of an analysis including a low number of cases, as long as an appropriate stratification correction is used. Our findings provide helpful guidelines for many researchers working on rare variant association studies.

## Introduction

Genetic association studies focusing on rare variants have become a popular approach to analyzing rare and common diseases. The advent of next-generation sequencing (NGS) and the development of new statistical approaches have rendered possible the comprehensive investigation of rare genetic variants, overcoming the shortcomings of classical genome-wide association studies (GWAS) [1, 2]. The main methods for testing rare variants for association do not test single variants against a phenotype, as in GWAS, but generally use an aggregation strategy within a genetic unit, usually a gene. These gene-based tests can be divided into two main categories: burden and variance-component tests [1-4]. Population stratification occurs when study subjects, usually cases and controls, are recruited from genetically heterogeneous populations. This problem is well known in association studies with common variants, causing an inflation of the type I error rate and reducing power. Several statistical approaches can be used to account for population stratification in GWAS. The most widely used are based on Principal Components (PC) analysis [5, 6] and Linear Mixed Models (LMM) [7-10].

Population stratification also affects association studies including rare variants [11-13]. However, it remains unclear whether the same correction methods can be applied to rare variant association studies [12, 14], particularly as rare and common variants may induce different types of population structure [12, 15]. Many studies have investigated the bias introduced by population stratification in the analysis of rare variants and have highlighted the need for corrective approaches to obtain meaningful results [12, 16, 17]. The performance of the correction method depends on the study setting and the method used to analyze the variants [11, 12, 18-21]. PC has been widely investigated [5, 6, 22-25] and shown to yield satisfactory correction at large geographic scales, but not at finer scales [20]. LMM have also been studied [19, 26] and shown to account for stratification well if variance-component approaches are used to test for association [19]. Most of these studies used simulated genetic data that did not completely reproduce the complexity of real exome sequences, and limited types of population structures. In addition, they used large numbers of cases (*e.g*. generally more than 500), which may not always be possible in practice, particularly in studies focusing on rare diseases.

We aimed at addressing such limitations of classical comparative studies with the comprehensive evaluation study proposed in this article. We investigated the main correction methods for rare variant association studies in the context of limited sample sizes, as in studies of rare disorders. For an accurate assessment of the different approaches, we used real NGS data from two sources: 1000 Genomes data [27] and our in-house cohort, with data for > 5,000 exomes [28]. We focused on two population structure scenarios: within-continent stratification (recent separation) and between-continent stratification (ancient separation). We also considered different sample sizes, including situations with as few as 50 cases, which have, to our knowledge, never been extensively investigated in this manner. We focused on a classic genetic model for a rare disease with a phenotype driven by rare deleterious variants well suited for a burden test, such as the cohort allelic sums test (CAST) [3]. We tested two classical correction methods, PC and LMM, a promising novel correction method called adapted local permutations (LocPerm) [29] and considered an uncorrected CAST-like test as a reference. Our global objective here is to provide useful practical insight into how best to account for population stratification in rare variant association studies.

## Materials and methods

### Simulation study

#### Exome data

For a realistic comparison of the correction approaches, we used two real exome datasets rather than program-based simulated exomes. Simulated data tend to provide erroneous site frequency spectra or LD structures [30]. The first dataset used was our HGID (Human Genetic of Infectious Diseases) database, containing 3,104 samples of in-house WES data generated with the SureSelect Human All Exon V4+UTRs exome capture kit (*https://agilent.com*). All study participants provided written informed consent for the use of their DNA in studies aiming to identify genetic risk variants for disease. IRB approval was obtained from The Rockefeller University and Necker Hospital for Sick Children, along with a number of collaborating institutions. The second dataset used was the 2,504 whole genomes from 1000 Genomes phase 3 (*http://www.internationalgenome.org/*) reduced with the same capture kit. We merged all the exomes from these two databases into a single large dataset before selecting samples. We performed quality control, retaining only coding variants with a depth of coverage (DP) > 8, a genotype quality (GQ) > 20, a minor read ratio (MRR) > 0.2 and call-rate > 95% [31]. We then excluded all related individuals based on the kinship coefficient (King’s kinship 2K > 0.1875) [32, 33], resulting in a final set of 4,887 unrelated samples. From these samples, we created two types of samples, as comparable as possible to those used in practice in association studies. The first sample, the “European” sample, consisted of samples from patients of European ancestry, and was used to assess stratification at the continental level. The second, the “Worldwide” sample, consisted of samples from European individuals together with North-African, Middle-Eastern, and South-Asian samples, for the assessment of intercontinental stratification.

#### European sample

We selected samples from individuals of European ancestry based on a reference sample and genetic distance. We first picked a European sample (sample HG00146 from the GBR population of 1000 Genomes, Figure 1A) and calculated its genetic distance to all other samples in the combined dataset. We used a Euclidean distance based on the first 10 PCs: the distance between individuals *i* and *j* is calculated as 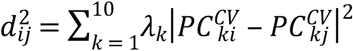, where ***PC***^***CV***^ is the matrix of principal components calculated on common variants and *λ*_*k*_ is the eigenvalue corresponding to the *k*-th principal component 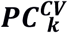. We considered that a sample could be “European” if its distance to the reference sample was below a certain threshold. This threshold was empirically chosen to ensure that all individuals of known European ancestry from the 1000 Genomes and our in-house HGID cohorts were included. The final sample consisted of 1,523 individuals, and included all the European samples from 1000 Genomes. We empirically separated the samples into three groups on the basis of ancestry (Figure 1B): Northern ancestry (including principally the FIN samples from 1000 Genomes), Middle-Europe ancestry (including the CEU and GBR samples from 1000 Genomes) and Southern ancestry (including the TSI and IBS samples from 1000 Genomes). The sample size for each subpopulation is shown in Table S1. After removal of the 102,219 private variants, the final sample contained 328,989 biallelic SNPs (Table S2).

**Figure 1.**
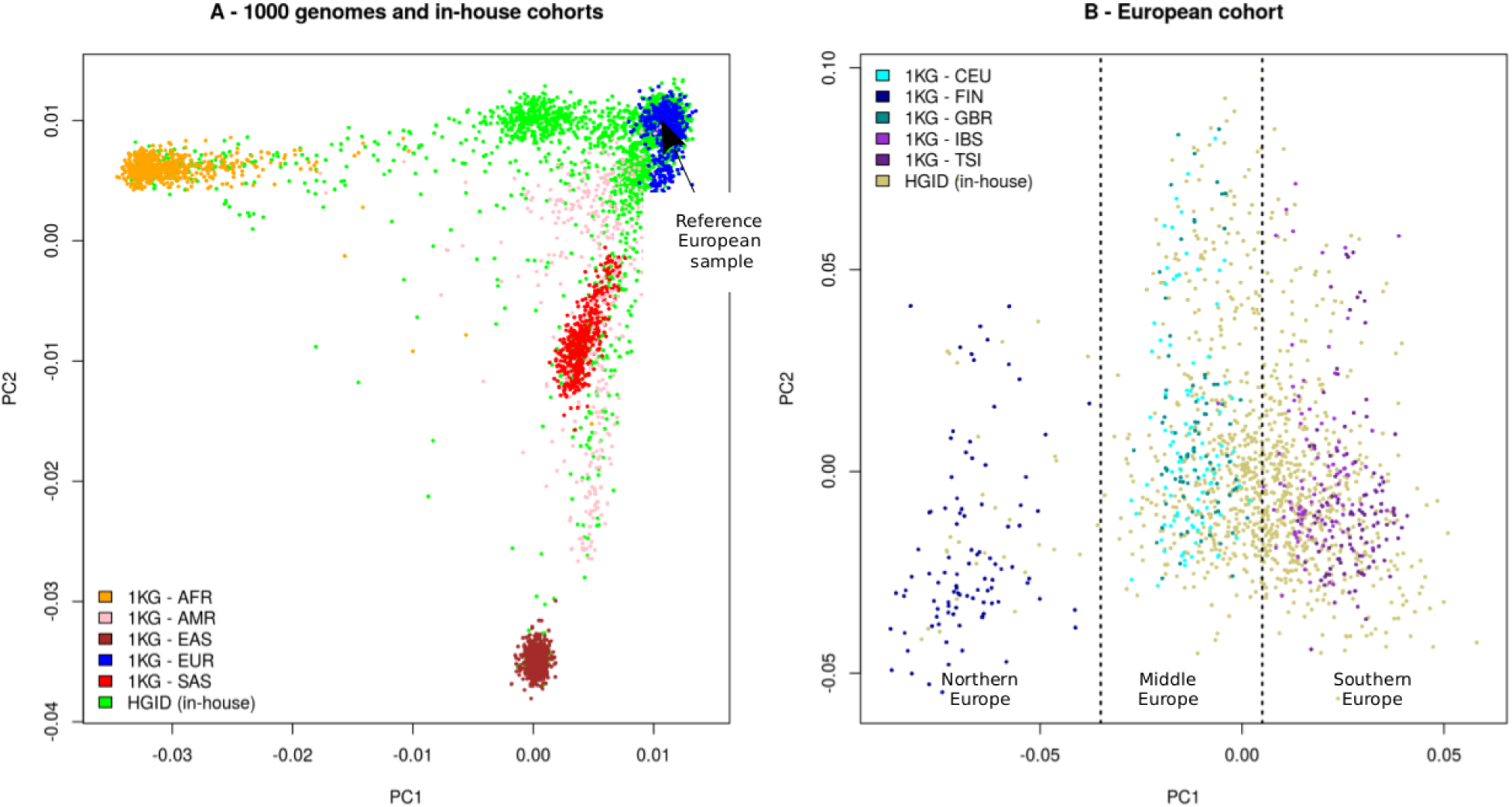
Graphical representation of the European sample. (A) PCA plots of the 4,887 samples comprising the 3,104 samples from our in-house cohort HGID and 1000 genomes (1KG) individuals including African (AFR), Ad Mixed American (AMR), East-Asian (EAS), European (EUR) and South-Asian (SAS). Common variants were used to produce these plots. The European reference individual is singled out. (B) 1,523 individuals with European ancestry selected. The dashed vertical lines correspond to empirical separations between Northern (n=127 including 1KG FIN and HGID samples), Middle (n=651 including 1KG CEU and GBR and HGID samples), and South European ancestry (n=745 including 1KG TSI and IBS and HGID samples).

#### Worldwide sample

The Worldwide sample was created in a similar manner. We selected four different reference samples of European (sample HG00146 from the GBR population of 1000 Genomes), South-Asian (sample NA20847 from the GIH population of 1000 Genomes), Middle-Eastern and North-African (samples from our in-house sample with a reported and verified Middle-Eastern or North-African ancestry) ancestry (Figure 2A). The genetic distances between each sample and the four reference samples were calculated as previously described. Thresholds were applied such that each sample with a reported ancestry of interest was assigned to the correct population and there was no overlap between the subpopulations (Figure 2B). The final Worldwide sample included 1,967 individuals separated into four subpopulations (Table S1). Note that all the European samples of this sample were also present in the European sample. This sample contained 483,762 biallelic SNPs after removal of the 132,565 private variants (Table S2).

**Figure 2.**
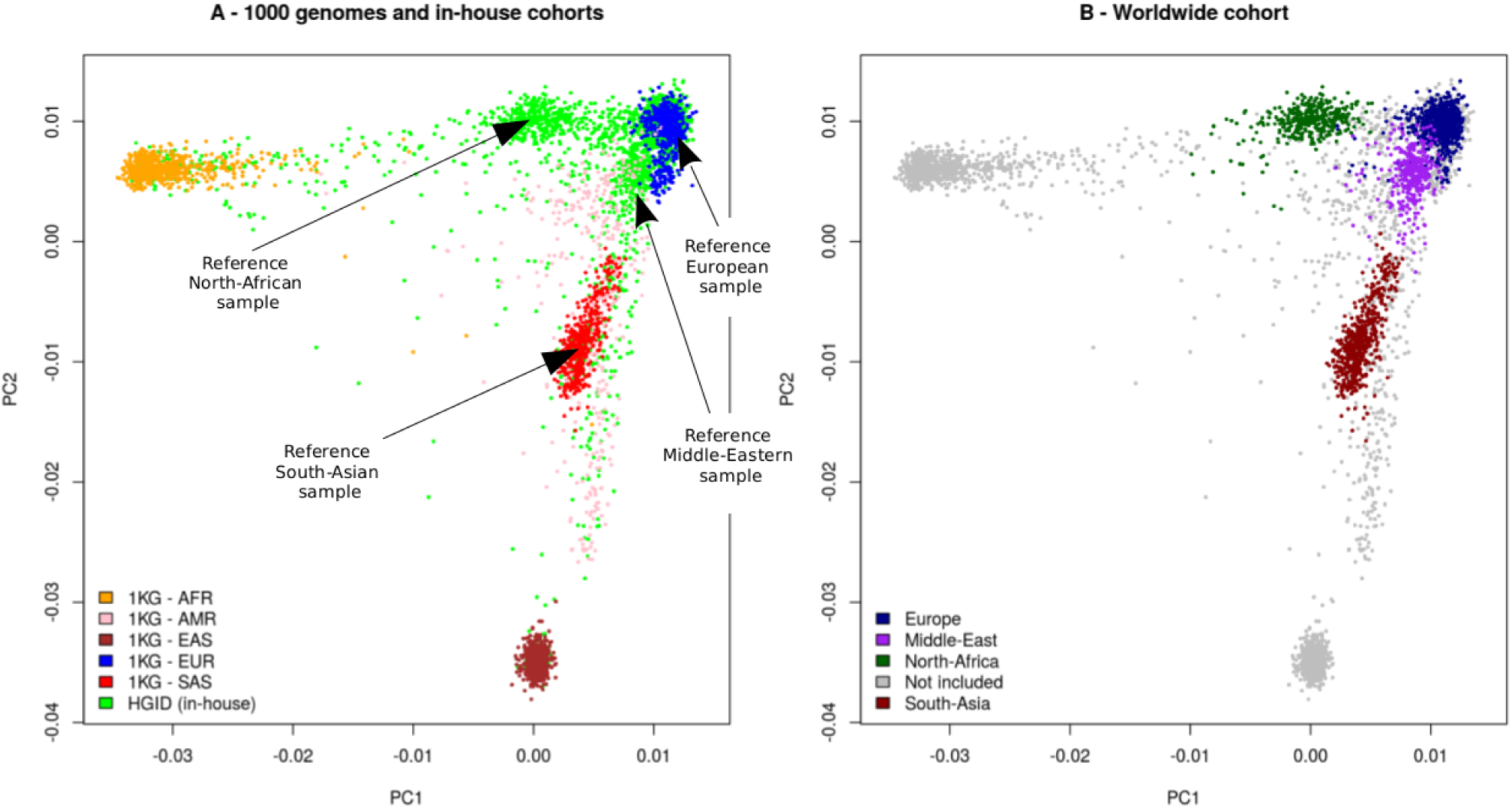
Graphical representation of the Worldwide sample. (A) PCA plots of the 4,887 samples comprising the 3,104 samples from our in-house cohort HGID and 1000 genomes (1KG) individuals including African (AFR), Ad Mixed American (AMR), East-Asian (EAS), European (EUR) and South-Asian (SAS). Common variants were used to produce these plots. Reference individuals are singled out. (B) The selected 1,967 individuals with European (n=700), Middle-Eastern (n=543), North-African (n=359) and South-Asian (n=365) ancestries are colored. The remaining individuals are left in grey.

#### Stratification scenarios

We first assessed the various correction approaches on case/control samples with large sample sizes (*i.e* with the whole European or Worldwide sample). We used the same three stratification scenarios for both samples. In each scenario, we considered a fixed proportion of 15% cases and 85% controls. Thus, in all our scenarios, the case/control ratio was unbalanced, as is often the case in practice. Comparison studies generally consider balanced scenarios with large numbers of cases and controls, corresponding to the ideal situation for most correction approaches, and their performance in more realistic conditions may therefore be overestimated. We considered a first scenario without stratification (No PS), in which we randomly selected 15% of the samples in each subpopulation as cases, the rest being used as controls. The second scenario corresponded to moderate stratification (Moderate PS), with the cases selected mostly from certain subpopulations. The third scenario was an extreme situation (High PS), in which all the cases were selected from a single subpopulation. The distribution of cases for the European and the Worldwide samples is shown, for each scenario, in Table 1.

**Table 1:**
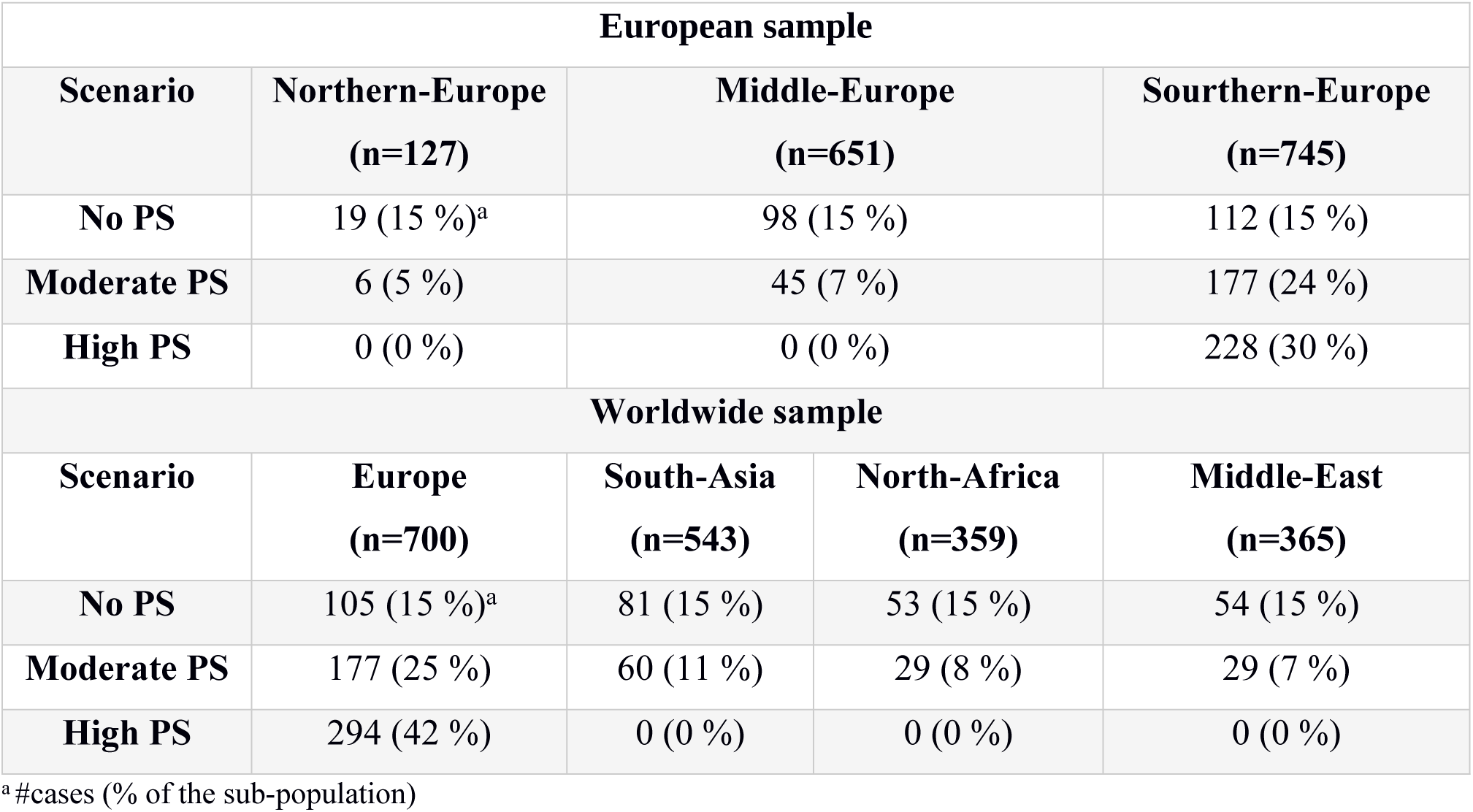
Distribution of the cases in the sub-populations of the European and the Worldwide samples for the different population stratification (PS) scenarios.

In practice, the samples used in rare variant association studies are frequently not very large. This is particularly true for rare diseases, for which only small numbers of cases are available. Case numbers may also be small as a consequence of the WES cost. The usual analysis strategy involves matching the controls to the cases. One key question is whether the addition of unmatched controls could increase the power of the analysis when population stratification is taken into account properly. Such controls are now available in large cohorts, such as the 1000 Genomes (Genomes Project, Auton (27)), UK10K [34], and UK Biobank [35] cohorts. We decided to investigate such strategies, by considering several scenarios with 50 cases and various numbers of controls of similar or different ancestries (Table 2). We considered three possible types of cases: 50 cases from the rather homogeneous Southern-Europe subpopulation (50SE), 50 cases from the more heterogeneous whole European population (50E) and 50 cases selected Worldwide (50W). Four types of controls were considered: 100 controls from the same population as the cases (100SE, 100E, 100W), 1000 controls from the total European sample (1000E), 1000 controls randomly chosen from the total Worldwide sample (1000W) and 2000 controls randomly chosen from the total Worldwide sample (2000W).

**Table 2:** Stratification scenarios for the small size study. The first 4 scenarios correspond to cases from the Southern-Europe sub-population (SE), the following 4 scenarios to cases from whole European sample (E) and the final 4 to cases from the Worldwide population (W). Controls are randomly drawn among the Southern-European, European or Worldwide populations.

| Scenario | Cases | Controls |
| --- | --- | --- |
| **50SE-100SE** | 50 from Southern-Europe | 100 from Southern-Europe |
| **50SE-1000E** | 50 from Southern-Europe | 1000 from all Europe |
| **50SE-1000W** | 50 from Southern-Europe | 1000 Worldwide |
| **50SE-2000W** | 50 from Southern-Europe | 2000 Worldwide |
| **50E-100E** | 50 from all Europe | 100 from all Europe |
| **50E-1000E** | 50 from all Europe | 1000 from all Europe |
| **50E-1000W** | 50 from all Europe | 1000 Worldwide |
| **50E-2000W** | 50 from all Europe | 2000 Worldwide |
| **50W-100W** | 50 Worldwide | 100 Worldwide |
| **50W-1000E** | 50 Worldwide | 1000 from all Europe |
| **50W-1000W** | 50 Worldwide | 1000 Worldwide |
| **50W-2000W** | 50 Worldwide | 2000 Worldwide |

#### Type I error rate evaluation

For each type of sample and stratification scenario, the type I error rate was estimated under the null hypothesis of no association between a gene and the phenotype (*H*_*0*_). We therefore simulated phenotypes, for the large sample, by randomly assigning the case and control states according to the stratification proportions provided in Table 1, respecting a fixed proportion of cases of 15%. Each protein-coding gene was then tested for association with the phenotype by the various statistical approaches described in the Statistical methods section. The rare variants included in these tests were biallelic variants with a *MAF*⩽5% in the sample analyzed. We included only genes with at least 10 rare variant carriers, resulting in 17,619 genes being studied in the European sample, and 17,854 genes in the Worldwide sample. A similar simulation process was applied to the small samples, according to the proportions of cases and controls described in Table 2. In these scenarios, the number of genes with at least 10 mutation carriers retained depended on sample size (Table S3). This procedure was repeated 10 times for each sample, to account for sampling variation. The type I error rate at the nominal level *α* was evaluated by assessing the quantity 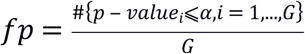 where *G* is the total number of genes tested. We decided to provide an adjusted prediction interval (PI), accounting for the large number of methods investigated, with the type I error rate as suggested in previous studies [19]. The bounds of this interval are 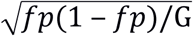 where *Z*_0.975/#(*methods*)_ replaces the usual 97.5 percentile of the normal distribution *Z*_0.975_ after adjustment for the number of methods investigated. An approach was considered to provide a good correction if its type I error rate was found within this interval.

#### Power studies

Power was estimated under the alternative hypothesis of an association between a gene and the phenotype (*H*_*1*_). We selected a subset of 10 genes for the power analysis. All these genes had a cumulative frequency of rare variants (*i.e* with *MAF*⩽5%) of ∼10% (*i.e*. ∼20% of carriers) and at least 10 mutation carriers. In addition, we considered ∼50% of the rare variants of each gene to be causal, with the same direction of effect, and used the presence of at least one of these variants to define the binary genetic score described in the Statistical method section. This implies that there was no cumulative effect of carrying several causal variants, and that the relative risk is defined at the gene level. Table S4 provides details of the 10 genes selected and their causal variants for the European and Worldwide samples. For each gene tested, a phenotype was simulated, using a binomial distribution and penetrance as parameters. For each stratification scenario, penetrance was calculated from the proportion of cases and controls, the frequency of carriers, and the relative risk (*RR=1,2,3,4*). An example is presented in Table S5 for the first gene tested. Tests of association between the genes and the simulated phenotypes were performed 500 times per gene, and power was estimated by evaluating the same quantity as for the type I error rate averaged over the 10 genes and the 500 replicates.

### Statistical methods

#### Association test

Let us now consider an association study including *n* individuals. The binary phenotype is denoted ***Y*** *= (y*_*1*_, …, *y*_*n*_), where *y*_*i*_ is the status of individual *i* coded 0 (healthy) or 1 (affected). We call ***X*** *= (x*_*ij*_) _*i=1*…*n, j=1*…*p*_ the *n x p* genotype matrix for *n* individuals and *p* markers. Each term *x*_*ij*_ corresponds to the genotype of sample *i* at marker *j* and is coded 0, 1 or 2 according to the number of minor alleles. We also introduce the normalized genotype matrix 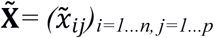, where each term is 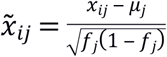 with *μ*_*j*_ the column mean and *f*_*j*_ the observed allele frequency of each marker.

Several routine statistical tests are available for assessing the association between rare variants and a phenotype. Considering our focus on a small number of cases with phenotypes driven by the presence of at least one causal variant, the most appropriate approach is that based on the CAST method [3]. This approach collapses variants into a single genetic score that takes a value of 0 if there are no rare variants in the region or 1 if there is at least one variant. Considering a given genetic region *g*, in our case a gene, the score for this region is denoted ***Z***_***g***_ *= (z*_*g1*_, …, *z*_*gn*_), where *z*_*gi*_ = *I* (at least one rare variant in the region *g* for individual *i*), *I()* being the indicator function.

The corresponding association test can be expressed in a logistic regression framework.

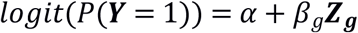

Where *α* and *β*_*g*_ are the model parameters for the intercept and the genetic score. Under the null hypothesis of no association {*β*_*g*_ = 0} the likelihood ratio test (LRT) statistics follow a 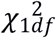 distribution.

#### Genetic similarity

Certain methods, including PC and LMM, account for population stratification by using a large number of single-nucleotide polymorphisms (SNPs) to derive genetic similarity matrices (also called relatedness matrices). Considering a set *H* of *p*_*H*_ SNPs, a normalized similarity matrix 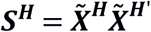 can be derived, where 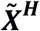 is the normalized genotype matrix reduced to the markers of set *H*. Each term *s*_*ik, i=1*…*n, k=1*…*n*_ represents the genetic similarity between samples *i* and *k* based on the SNPs of set *H*.

With whole-exome sequencing (WES) data, a broad range of SNPs are now available, and it is usual to separate them into categories based on their minor allele frequencies (MAFs) [18, 19, 24]. We will consider four categories of variants, based on the MAFs calculated for the total sample: rare variants (RVs; 0% < *MAF* < 1%), low-frequency variants (LFVs; 1%⩽*MAF* < 5%), common variants (CVs; *MAF*⩾5%) and all variants (ALLVs; the union of RVs, LFVs and CVs). We excluded private variants from these sets of variants, because their sparse distribution tends to have a strong influence on the calculation of similarity matrices. We also pruned all these sets to remove variants with a pairwise r^2^ < 0.2, to reduce the effect of linkage disequilibrium. We investigated the effect of using these different sets of SNPs *H* ∈ {*RVs,LFVs,CVs,ALLVs*} to derive PC-based or LMM corrections.

#### Principal component (PC) approach

PC analysis creates new variables from SNP data, the principal components, corresponding to axes of genetic variation. These variables can be included, as covariates, in a statistical model, such as the one described above to adjust for population stratification. Principal components 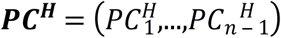 are based on a given set of SNPs *H* and are derived from the singular vector decomposition of the normalized similarity matrix ***S***^***H***^. After adjustment for the first *m* principal components, the corresponding logistic model becomes:

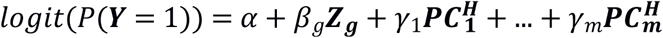

where *γ*_1,_…,*γ*_*m*_ are new model parameters for the PCs.

Under the null hypothesis of no association {*β*_*g*_ = 0}, the LRT statistics follow a 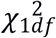 distribution. We investigated correction based on the first 3, 5, 10 or 50 PCs, calculated on the four possible sets of variants, RVs, CVs, LFVs and ALLVs. In the following, we use a notation such that PC3_CV_, for example, indicates that the first three PCs based on common variants were used.

#### Linear mixed models (LMM)

Linear mixed models were initially developed to alleviate the effect of familial relatedness in association analyses, and have also been used to correct for population stratification in GWAS. This regressive approach considers both fixed and random effects and uses a genetic similarity matrix to improve estimation of the parameters of interest. Using the previous CAST regression framework, the LMM model becomes:

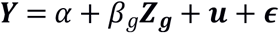

where ***u*** ∼ *MVN*(0,*τ****SH***) is a vector of random effects based on the similarity matrix ***S***^***H***^ and an additional variance parameter *τ*. Under the null hypothesis of no association {*β*_*g*_ = 0}, the LRT statistics follow a 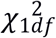 distribution. We focus here on LMM based on the relatedness matrices constructed with the four sets of variants previously described, and with for instance the notation LMM_CV_ indicating that common variants were used.

#### Adapted local permutations (LocPerm)

Permutation strategies have been designed to derive *p-*values when the ‘true’ null distribution of the test statistic *T*_*0*_ is unknown [36]. This is the case for population stratification, which creates a bias that cannot be numerically derived. The rationale behind permutation procedures is to simulate several test statistics *(T*_*1*_, …, *T*_*B*_) under the null hypothesis, to derive an approximated distribution as close as possible to the unknown true null distribution, and to use these statistics to estimate a *p-*value. With the classical permutation approach, the simulation of test statistics under *H*_*0*_ is achieved by randomly resampling phenotypes (*i.e* exchanging them between individuals). Adapted local permutations are based on the observation that, in the presence of population structure, not all phenotypes are exchangeable [29]. A given sample has a higher chance of sharing its phenotype with another sample of the same ancestry. The principle is, therefore, to establish, for each sample, a neighborhood, *i.e* a set of samples between which it is reasonable to exchange phenotypes. These neighborhoods are established according to a genetic distance derived from the first 10 PCs:

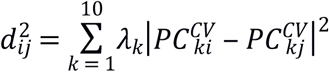

where ***PC***^***CV***^ is the matrix of principal components calculated on the set of common variants and *λ*_*k*_ is the eigenvalue corresponding to the *k*-th principal component 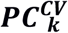. This distance is used to create a neighborhood of 30 individuals around each sample [29]. Permutations can then be performed for each sample, within its neighborhood.

A straightforward empirical way to derive a *p-*value for the permutation test is to assess the quantity *pv* = #{*T*_*i*_⩾*T*_0_}/*B* where # is the cardinal function and *B* is the number of permutations. This method is dependent on the number of permutations computed, and a large number of permutations is required for the accurate estimation of small *p*-values. Mullaert et al. proposed an alternative semi-parametric approach, in which a limited number of resampled statistics are used to estimate the mean (m) and standard deviation (σ) of the test statistic under *H*_*0*_. The previously described CAST-like LRT statistics are used, through their square roots with a sign attributed according to the direction of the effect, 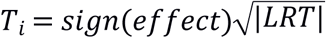, to estimate the N(m, σ^2^) distribution parameters and then calculate the *p-*value. We evaluated both the semi-parametric approach using 500 local permutations and the full empiric approach using 5000 local permutations. These two approaches yielded very similar results. We therefore present here only the results for the semi-empiric approach.

#### Implementation of the simulations and methods

We used R software (*https://www.R-project.org/*) to code the comparison pipeline and implement the logistic and permutation models. Principal components and similarity matrices were obtained with Plink2 software (*https://www.cog-genomics.org/plink/2.0/*), and GEMMA was used for the LMM method [10, 37].

## Results

### Large study size

The results of the simulation study under the null hypothesis for the European sample of 1,523 individuals are presented in Table 3 (for α=0.001) and Table S6 (for α=0.01). In the absence of stratification, the four methods had correct type I error rates, within the 95% PI bounds (Table 3A, Table S6A). This was the case for PC3 and LMM, regardless of the type of variant considered. In the presence of moderate stratification (Table 3B, Table S6B), the unadjusted CAST approach displayed the expected inflation of type I error rate (0.00163 at α=0.001). The PC3 method corrected properly regardless of the type of variant at α=0.001, but a slight inflation of type I error was observed for RVs and LFVs at α=0.01. The use of LMM led to an inflation of type I error rates at α=0.001, unless all variants were considered, which gave rates within the 95% PI at α=0.01. LocPerm had a correct type I error rate at both α levels. In the presence of strong stratification (Table 3C, Table S6C), the unadjusted CAST method gave a strong inflation of type I error rate, to 0.00359 at α=0.001. The PC and LMM approaches also led to inflated type I errors (between 0.00133 and 0.00175 at α=0.001), the lowest level of inflation being observed when CVs or all variants were considered. For the PC approach, increasing the number of PCs did not improve the correction, consistent with previous findings reported by Persyn et al. (2018). The use of 50 PCs resulted in an inflation of type I error whatever the level of stratification, probably due to an overadjustment of the regression model (Table S7). Thus, in the presence of strong population structure, classical methods were unable to handle the stratification properly. The adapted local permutations approach was the only method able to correct for stratification in this scenario, with a slightly conservative result of 0.00863 at α=0.01 (Table S6C).

**Table 3:** Type I error rates of the different approaches for the large size European sample. The nominal level alpha considered is *α* = 0.001 and the corresponding 95%PI adjusted for the 10 methods is [0,00079-0,00121]. Type I error rates under the lower bound of the 95%PI are displayed in italic and above the upper bound of the 95%PI in bold.

| A - No Stratification |  |  |  |  |
| --- | --- | --- | --- | --- |
|  | CAST | PC3 | LMM | LocPerm |
| RVs | 0.00106 | 0.00108 | 0.00118 | 0.00082 |
| LFVs |  | 0.0011 | 0.00119 |  |
| CVs |  | 0.00104 | 0.00118 |  |
| ALLVs |  | 0.00108 | 0.00116 |  |
| B – Moderate Stratification |  |  |  |  |
|  | CAST | PC3 | LMM | LocPerm |
| RVs | 0.00163 | 0.00117 | 0.00141 | 0.00095 |
| LFVs |  | 0.00101 | 0.00125 |  |
| CVs |  | 0.001 | 0.00124 |  |
| ALLVs |  | 0.00102 | 0.00117 |  |
| C – High Stratification |  |  |  |  |
|  | CAST | PC3 | LMM | LocPerm |
| RVs | 0.00359 | 0.00157 | 0.00175 | 0.00087 |
| LFVs |  | 0.00137 | 0.00176 |  |
| CVs |  | 0.00136 | 0.00161 |  |
| ALLVs |  | 0.00133 | 0.00145 |  |

The results of the simulation study under *H*_*0*_ for the Worldwide sample of 1,967 individuals are presented in Table 4 (for α=0.001) and Table S8 (for α=0.01). In the absence of stratification, none of the main approaches had a significantly inflated type I error rate (Table 4A and Table S8A). At *α* =0.01, LMM corrections were slightly conservative. The presence of moderate or strong stratification led to extremely inflated type I errors at α=0.001 for the unadjusted CAST approach, with values of 0.00681 and 0.137, respectively. For PC3 and LMM, a satisfactory correction was obtained at α=0.001 with CVs, whereas, at α=0.01, PC gave a slight inflation of type I error and LMM results were slightly conservative. The three other types of variants could not properly account for stratification for PC3 and LMM. Increasing the number of PCs did not improve the results obtained with PC3 (Table S9) for the Worldwide sample. LocPerm maintained a correct type I error rate in both scenarios, with values of 0.00096 and 0.00113 at α=0.001 for moderate and strong stratification, respectively. Overall, the analyses under the null hypothesis within the European and Worldwide samples showed that accounting for stratification was generally more difficult with a continental structure than with a worldwide structure. PC3 and LMM based on CVs were capable of maintaining a correct type I error rate in most of the situations considered, with the exception of high levels of stratification in Europe, and LocPerm correctly accounted for stratification in all the situations considered.

**Table 4:**
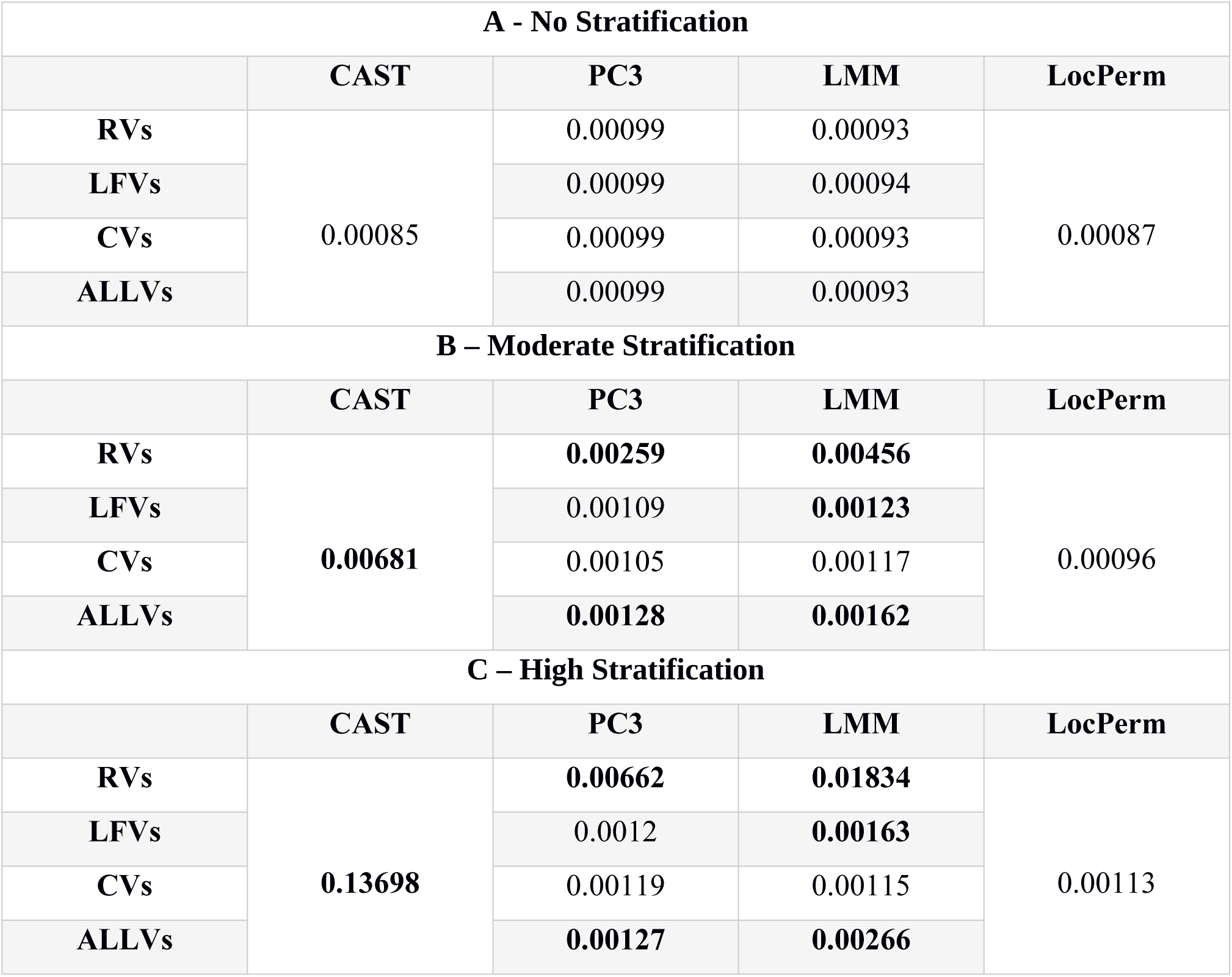
Type I error rates of the different approaches for the large size Worldwide sample. The nominal level alpha considered is *α* = 0.001 and the corresponding 95%PI adjusted for the 10 methods is [0,00079-0,00121]. Type I error rates under the lower bound of the 95%PI are displayed in italic and above the upper bound of the 95%PI in bold.

| A - No Stratification |  |  |  |  |
| --- | --- | --- | --- | --- |
|  | CAST | PC3 | LMM | LocPerm |
| RVs | 0.00085 | 0.00099 | 0.00093 | 0.00087 |
| LFVs |  | 0.00099 | 0.00094 |  |
| CVs |  | 0.00099 | 0.00093 |  |
| ALLVs |  | 0.00099 | 0.00093 |  |
| B – Moderate Stratification |  |  |  |  |
|  | CAST | PC3 | LMM | LocPerm |
| RVs | 0.00681 | 0.00259 | 0.00456 | 0.00096 |
| LFVs |  | 0.00109 | 0.00123 |  |
| CVs |  | 0.00105 | 0.00117 |  |
| ALLVs |  | 0.00128 | 0.00162 |  |
| C – High Stratification |  |  |  |  |
|  | CAST | PC3 | LMM | LocPerm |
| RVs | 0.13698 | 0.00662 | 0.01834 | 0.00113 |
| LFVs |  | 0.0012 | 0.00163 |  |
| CVs |  | 0.00119 | 0.00115 |  |
| ALLVs |  | 0.00127 | 0.00266 |  |

With respect to the results of the simulation under *H*_*0*_, we focused the power studies on the methods providing satisfactory correction (*i.e*. PC3_CV_, LMM_CV_ and LocPerm), in addition to the unadjusted CAST. Only powers derived from a correct type I error rate under *H*_*0*_ are presented in the main text. Adjusted powers accounting for inflated type I error rates are provided in the Supplementary figures for information. The results of the power study for the European sample are presented in Figure 3 and Figure S1. In situations with no stratification or moderate stratification, all approaches had similar powers, of about 50% at α=0.001 for a relative risk of 3, for example (Figure 3). In the presence of strong stratification, only LocPerm was able to correct for confounding and to maintain power levels (Figure 3C). The adjusted powers (Figure S1) indicate that all three correction methods provide very similar powers when type I error is controlled. The results of the power study for the Worldwide sample are presented in Figure 4 and Figure S2. As for the European sample, all methods had similar powers in the absence of stratification or the presence of moderate stratification. In the presence of strong stratification, LocPerm was slightly less powerful than the other methods (Figure 4C) with for a RR of 3 at α=0.001, a power of 64% as opposed to the powers of 69 and 72% obtained for PC3_CV_, and LMM_CV_, respectively. It is also interesting to compare the power of each method, separately, between the different stratification scenarios (Figures S3 for the European sample and S4 for the Worldwide sample). Power was very similar for any given technique in the different stratification scenarios, indicating that the correction methods maintained the level of power observed in the absence of stratification.

**Figure 3.**
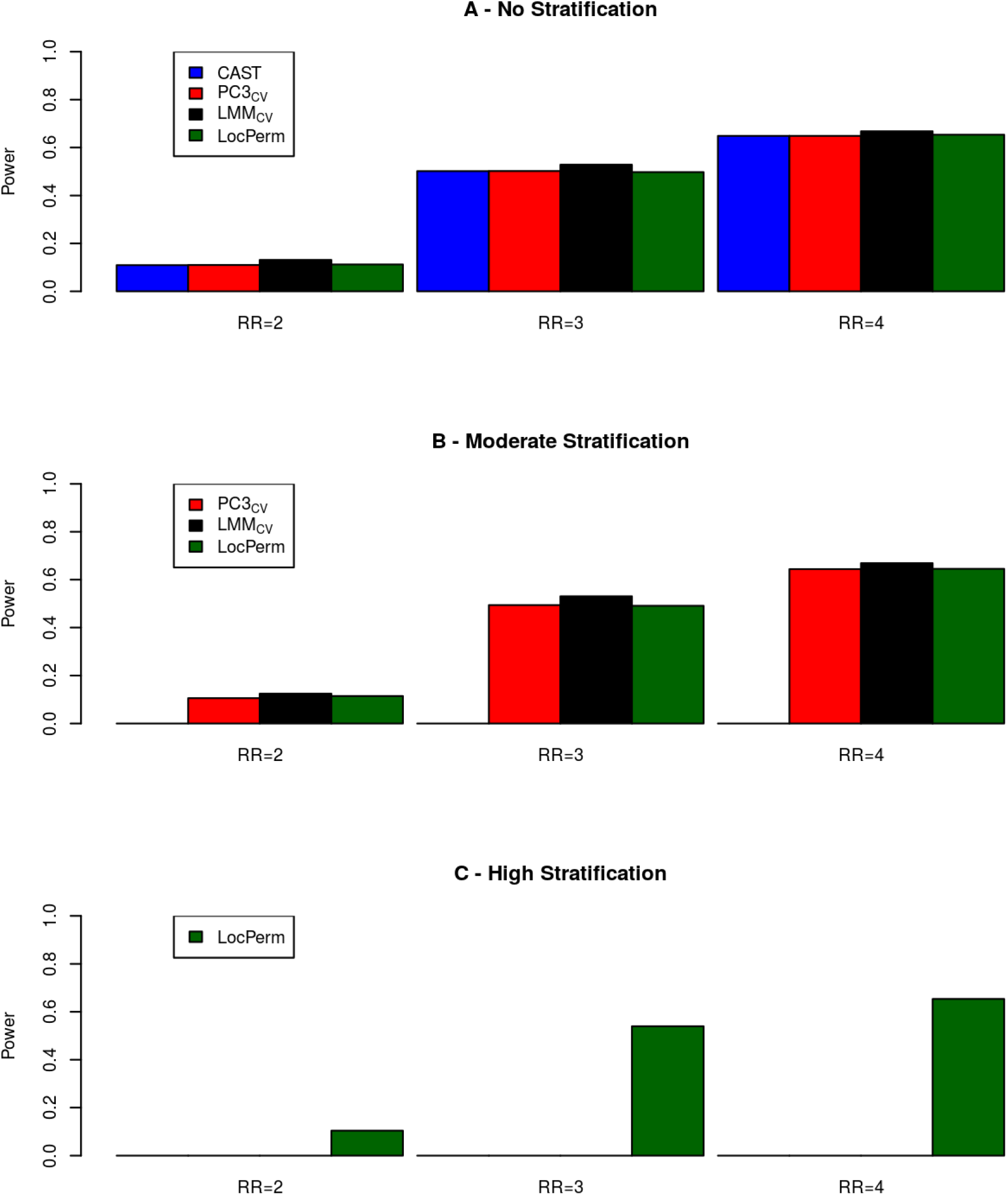
Histogram of powers for methods with a correct type I error rate for the large size European sample (n=1,523) at the level *α* = 0.001. (A) Without stratification. (B) With moderate stratification. (C) With high stratification. Relative risks considered vary from 2 to 4 on the x-axis.

**Figure 4.**
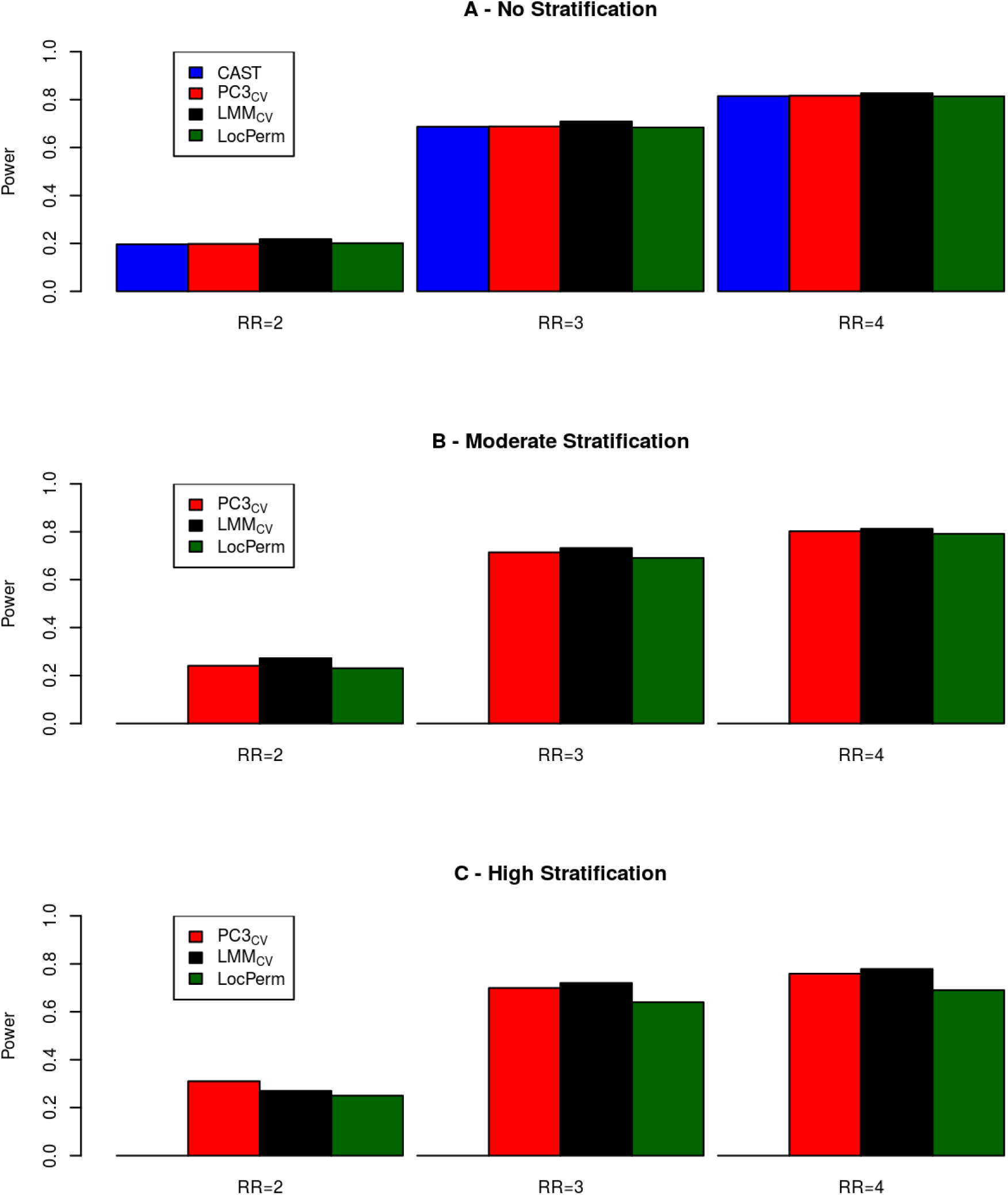
Histogram of powers for methods with a correct type I error rate for the large size Worldwide sample (n=1,967) at the level *α* = **0.001**. (A) Without stratification. (B) With moderate stratification. (C) With high stratification. Relative risks considered vary from 2 to 4 on the x-axis.

### Small study size

The results of the simulation study under the null hypothesis for a small sample size, based on 50 cases, are presented in Table 5 (for α=0.001) and Table S10 (for α=0.01). Only PC3_CV_, LMM_CV_ and LocPerm, which provided a satisfactory correction for stratification in the large sample study, were investigated for small sample sizes. In scenarios without stratification (i.e. controls and cases of the same origin), an inflation of type I errors was observed: 1) with PC3 (about 0.0015 at α=0.001) when the number of controls was low (100), and, to a lesser extent, with CAST (about 0.0012 at α=0.001), and 2) with LMM (about 0.002 at α=0.001) when the number of controls was high (1000 or 2000). In the presence of stratification (*i.e*. a large number of controls with an origin different from that of the cases), a strong inflation of type I error rates was observed for CAST. This was also the case for LMM_CV_, albeit to a lesser extent, particularly for stratification within Europe or when the cases came from the Worldwide sample and the controls from Europe only. Both PC3_CV_ and LocPerm provided correct type I error rates in all the scenarios considered with small numbers of cases and a large number of controls.

**Table 5:**
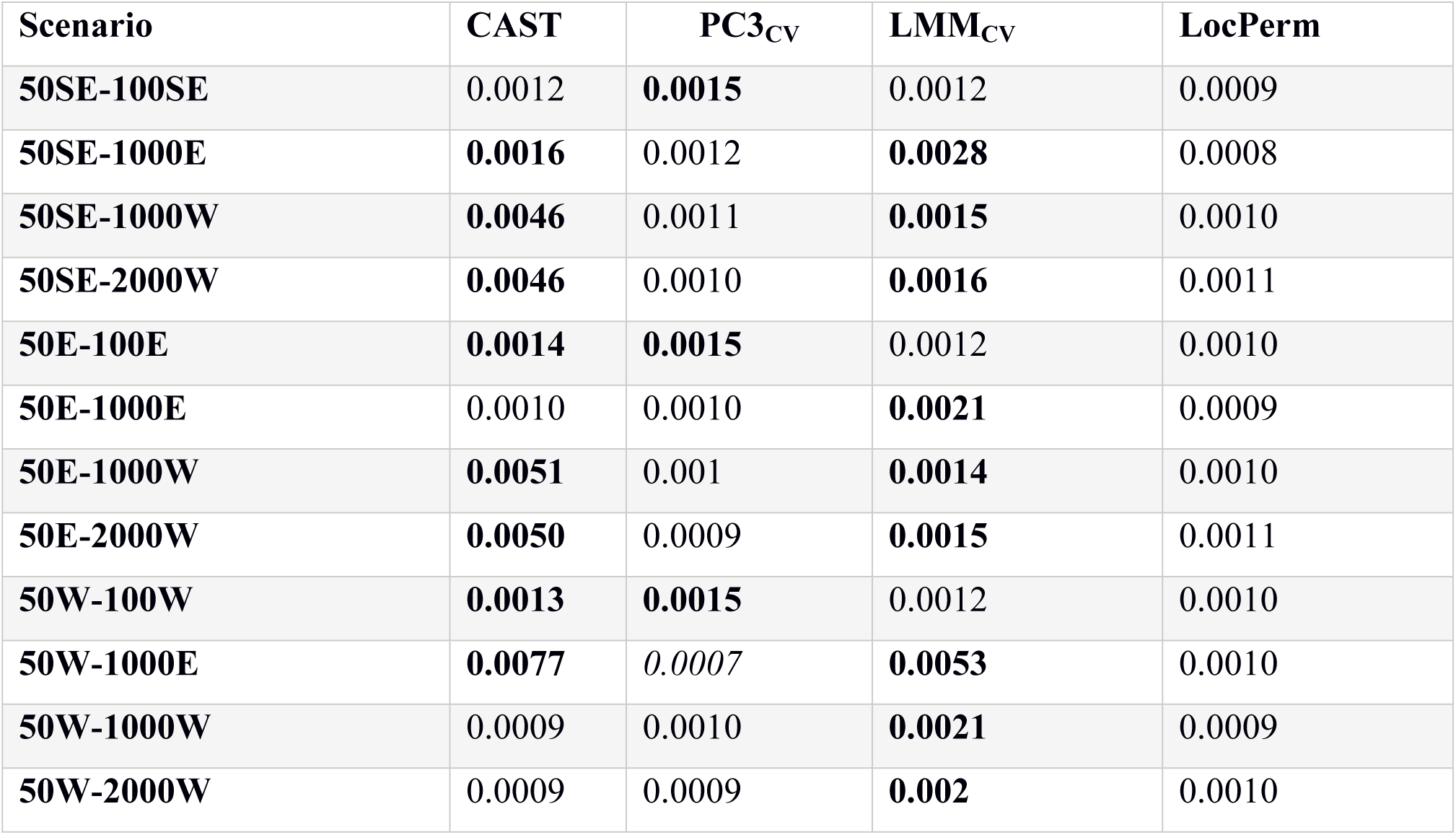
Type I error rates of the different approaches for the small size sample scenarios. The nominal level alpha considered is *α* = 0.001. Type I error rates under the lower bound of the 95%PI are displayed in italic and above the upper bound of the 95%PI in bold. STable 3 provides the adjusted 95%PI for the different number of genes tested in each scenario.

| Scenario | CAST | PC3<sub>CV</sub> | LMM<sub>CV</sub> | LocPerm |
| --- | --- | --- | --- | --- |
| 50SE-100SE | 0.0012 | **0.0015** | 0.0012 | 0.0009 |
| 50SE-1000E | **0.0016** | 0.0012 | **0.0028** | 0.0008 |
| 50SE-1000W | **0.0046** | 0.0011 | **0.0015** | 0.0010 |
| 50SE-2000W | **0.0046** | 0.0010 | **0.0016** | 0.0011 |
| 50E-100E | **0.0014** | **0.0015** | 0.0012 | 0.0010 |
| 50E-1000E | 0.0010 | 0.0010 | **0.0021** | 0.0009 |
| 50E-1000W | **0.0051** | 0.001 | **0.0014** | 0.0010 |
| 50E-2000W | **0.0050** | 0.0009 | **0.0015** | 0.0011 |
| 50W-100W | **0.0013** | **0.0015** | 0.0012 | 0.0010 |
| 50W-1000E | **0.0077** | *0.0007* | **0.0053** | 0.0010 |
| 50W-1000W | 0.0009 | 0.0010 | **0.0021** | 0.0009 |
| 50W-2000W | 0.0009 | 0.0009 | **0.002** | 0.0010 |

A power study was performed for PC3_CV_ and LocPerm with small numbers of cases (Figure 5). Both approaches gave a correct type I error rate and similar results, but power was slightly higher for LocPerm than for PC3 when the 50 cases came from Europe as a whole or from the Worldwide sample. For cases were from Southern Europe, considering 1000 controls from the whole of Europe gave a power twice that obtained when considering 100 controls of the same origin as the cases (Figure 5A). For example, for a RR of 4 and at α=0.001, the power increased from 15% to 34% under these conditions with LocPerm. A smaller increase was observed if 1000 controls from the Worldwide sample were used, increasing to a similar level with the use of 2000 Worldwide controls. When the cases were from anywhere in Europe, a similar increase in power was observed with 1000 European and with 1000 Worldwide controls, whereas the use of 2000 Worldwide resulted in no greater a power than the use of 1000 Worldwide controls. Finally, when the cases were selected from the Worldwide sample, the use of 1000 Worldwide controls gave a power almost double that achieved with 100 Worldwide controls, whereas the use of 1000 controls from Europe did not substantially increase the power. These results indicate that using a large panel of worldwide controls to increase sample size is a good strategy for increasing the power of a study while correcting for stratification with approaches such as PC3_CV_ or LocPerm.

**Figure 5.**
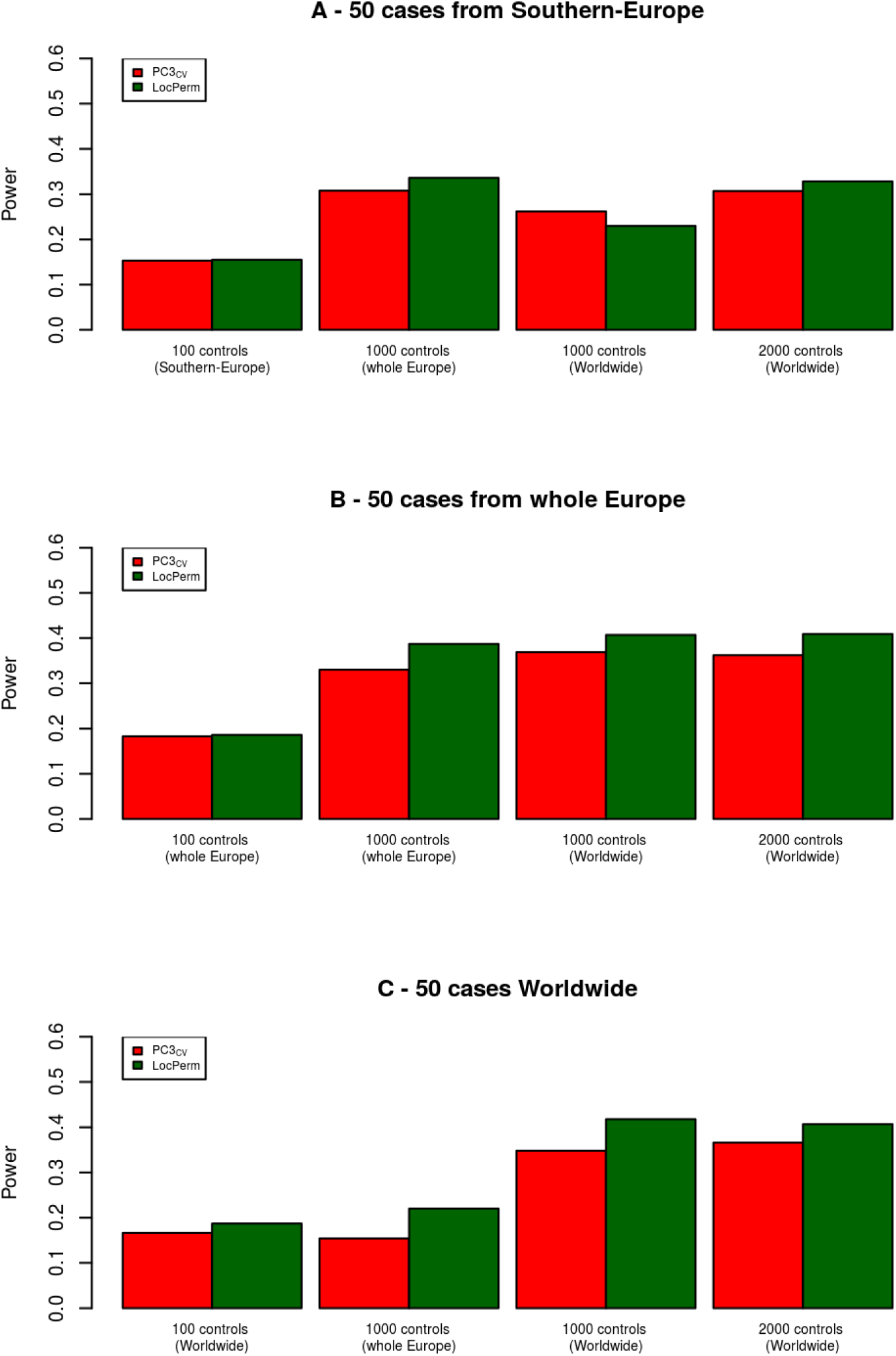
Power for methods with a correct type I error rate under H_0_ for the small size sample at the level *α* = 0.001. (A) Scenarios with 50 cases from Southern-Europe. (B)Scenarios with 50 cases from the whole Europe. (C) Scenarios with 50 cases from the Worldwide sample. The relative risk is fixed at 4.

### Computational considerations

We also assessed the computing time required for the different approaches. While the unadjusted CAST method does not imply the computation of any particular matrix, the same covariance matrix is necessary for PC3_CV_, LMM and LocPerm and additional specific permutation matrices are required for LocPerm only. We ran each method separately, CAST, PC3_CV_ and LocPerm with R, and LMM_CV_ with GEMMA, on the 1,523 individuals and the 17,619 genes of the European sample, under a hypothesis of no association. We broke down the runtime of each method into a pretreatment phase (covariance and permutation matrices) and a gene-testing phase (see Table S11). The pretreatment runtime was dependent only on the number of individuals (and the set of SNPs used for the calculations) and this part of the analysis was performed only once. The runtime of the gene-testing phase depended on the number of individuals and the number of genes tested, and could be repeated for different analyses (*e.g*. for different MAF thresholds). PC3_CV_ and LMM_CV_ had similar pretreatment times, markedly shorter than that for LocPerm, which also requires the calculation of permutation matrices. However, the need to calculate these matrices only once decreases the relative disadvantage of the LocPerm method. In terms of gene-testing time, LMM_CV_ was the fastest approach when used with GEMMA, but this may not be the case for other programs that have not been optimized. A comparison of the methods implemented with R showed that the adjustment on PCs and LocPerm took 1.4x and 2.5x longer, respectively, than the unadjusted test. These comparisons were run on a 64-bit Intel Xeon Linux machine with a CPU of 3.70 GHz and 64 GB of RAM.

## Discussion

We performed a large simulation study based on real exomes data to investigate the ability of several approaches (i.e. PCs, LMM and LocPerm) to account for population stratification in rare-variant association studies of a binary trait. In our simulation study, the efficiency of PCs and LMM to correct for population stratification was dependent on the type of variant used to derive the similarity matrices, the best performance being obtained with CVs. It was generally not possible to correct the stratification bias with RVs, even with the exclusion of private variants for the calculation of the matrices. Private variants have very sparse distributions, which may lead to difficulties in calculation, and their inclusion resulted in an even lower efficiency of correction for population structure (data not shown). Other studies evaluating different types of variants reached the same conclusions [24, 25] although one reported better performances for PC based on RVs [14]. However, this study was based on simulated NGS data, which may have led to an unrealistic rare variant distribution. Our results also indicate that CVs or ALLVs were the best sets of variants for the LMM approach applied to CAST, confirming the results of Luo et al. based on the SKAT test [19]. Variant selection remains an area in which there are perspectives for improving the corrections provided by strategies such as PC or LMM [12, 26], although the use of CVs appeared to be a good choice in most situations.

With the optimal set of variants, PC generally corrected for population stratification more efficiently than LMM. This is consistent with benefits of the PC approach over LMM observed in the presence of spatially confined confounders [38], which is often the case with rare variants. For large sample sizes, both PC and LMM controlled for stratification better at larger geographic scales than at finer scales. In small samples (50 cases and 100 controls), PC approaches gave inflated type I errors even in the absence of population stratification, as previously reported [18, 29, 39]. This inflation disappeared when the sample included additional controls, whatever their ethnic origin, even with a highly unbalanced case-control ratio. By contrast, the type I error of LMM was inflated in samples with highly unbalanced case-control ratios, whatever the level of population stratification, as previously noted in the context of GWAS [40]. Finally, the adapted local permutations procedure recently proposed by Mullaert et al. [29] gave very promising results, as it fully corrected for population stratification, regardless of the scale over which the stratification occurred, sample size and case-control ratio. When valid under H_0_, the three correction methods had similar powers. For a given setting, power was similar in the different stratification situations, indicating that the correction method could maintain the power it would have in the absence of stratification. These results are in partial agreement with several studies reporting a small loss of power for PC-adjusted logistic regression in the presence of stratification relative to an absence of stratification [13, 20].

We also investigated the specific situation in which only a very small number of cases are available, which is particularly relevant in the context of rare disorders. In this setting, we show that PC and LocPerm provide correct type I errors when the number of controls is large, regardless of the ethnic origin of the controls. In addition, the strategy of adding controls, even of worldwide origin, provided a substantial gain of power for PC and LocPerm when few cases were available. This is an important finding, highlighting the potential interest of using publicly available controls, such as those of the 1000G project, to increase the power of a study with a small sample size. We also investigated an additional scenario in which all cases were strictly from our in-house HGID cohort and the controls were obtained from both the HGID and 1000 Genomes cohorts (data not shown). This scenario gave identical results to those presented here, indicating that, even in the presence of heterogeneity in the types of exome data considered for cases and controls (*e.g*. in terms of kit or technology used), the conclusions drawn here still apply. Overall, these results validate a strategy of using additional external controls to increase the power of a study, provided that an efficient stratification correction approach is used.

We focused on the investigation of diseases caused by a few deleterious variants, for which the CAST-like approach is particularly appropriate. Additional studies are required to investigate more complex genetic models, such as the presence of both risk and protective variants of a given gene, for which other association tests, such as variant-component approaches, may be more appropriate. Different results can be expected, as the effect of population stratification differs between testing strategies [17, 20]. In addition, the novel LocPerm strategy has not been evaluated in combination with other association tests. In the situations we considered, our study highlighted several useful conclusions for rare variant association studies in the presence of stratification: 1) the key issue is to properly control type I errors as powers are comparable, 2) population stratification can be corrected by PC3_CV_ in most instances, unless there is a high degree of intracontinental stratification and a small sample size, 3) LocPerm proposes a satisfying correction in all instances, and 4) strategies based on the inclusion of a large number of additional controls (*e.g*. from publicly available databases) provide a substantial gain of power provided that stratification is controlled for correctly.

## Acknowledgments

We thank both branches of the Laboratory of Human Genetics of Infectious Diseases for helpful discussions and support. The Laboratory of Human Genetics of Infectious Diseases was supported in part by grants from the French National Agency for Research (ANR) under the “Investissement d’avenir” program (grant number ANR-10-IAHU-01), the TBPATHGEN project (ANR-14-CE14-0007-01), the MYCOPARADOX project (ANR-16-CE12-0023-01), the Integrative Biology of Emerging Infectious Diseases Laboratory of Excellence (grant number ANR-10-LABX-62-IBEID), the St. Giles Foundation, the National Center for Research Resources and the National Center for Advancing Sciences (NCATS), and the Rockefeller University.

## Supporting information

**Table S1. Distribution of the samples in the European and Worldwide sub-populations**

**Table S2. Distribution of the variants in the European and the Worldwide samples according to their MAFs as described in the Material and Methods section**.

**Table S3. Number of genes tested and 95%PI in each scenario of the small size sample study**. Prediction intervals are adjusted on the 4 methods tested.

**Table S4. Details of the genes selected for the power analysis in the European and the Worldwide samples**. Freq() indicates the cumulative frequency of the causal variants.

**Table S5. Example of penetrances used for the power estimation of the gene ADAMTS4 in the European sample**. F_0_ and F_1_ represent the penetrances for non-carriers and carriers considering a relative risk RR of 3, a total proportion of 15% of cases and proportions of carriers of 9% in Northern-Europe (NE) and Middle-Europe (ME) and 8% in Southern-Europe (SE). Within a given sample these penetrances are calculate by F_0_ = n_cases_/(n_non-carriers_+RR.n_carriers_) and F_1_=RR.F_0_

**Table S6. Type I error rates of the different approaches for the large size European sample**. The nominal level alpha considered is *α* = 0.01 and the corresponding 95%PI adjusted for the 10 methods is [0.00933-0.01067]. Type I error rates under the lower bound of the 95%PI are displayed in italic and above the upper bound of the 95%PI in bold.

**Table S7. Type I error rates of the PC approach with 3, 5, 10 or 50 PCs for the large size European sample**. The nominal level alpha considered is *α* = 0.001 and the corresponding 95%PI adjusted for the 16 methods is [0.00078-0.00122]. Type I error rates under the lower bound of the 95%PI are displayed in italic and above the upper bound of the 95%PI in bold.

**Table S8. Type I error rates of the different approaches for the large size Worldwide sample**. The nominal level alpha considered is *α* = 0.01 and the corresponding 95%PI adjusted for the 10 methods is [0.00933-0.01067]. Type I error rates under the lower bound of the 95%PI are displayed in italic and above the upper bound of the 95%PI in bold.

**Table S9. Type I error rates of the PC approach with 3, 5, 10 or 50 PCs for the large size Worldwide sample**. The nominal level alpha considered is *α* = 0.001 and the corresponding 95%PI adjusted for the 16 methods is [0.00078-0.00122]. Type I error rates under the lower bound of the 95%PI are displayed in italic and above the upper bound of the 95%PI in bold.

**Table S10. Type I error rates of the different approaches for the small size sample scenarios**. The nominal level alpha considered is *α* = 0.01. Type I error rates under the lower bound of the 95%PI are displayed in italic and above the upper bound of the 95%PI in bold.

Table S3 provides the adjusted 95%PI for the different number of genes tested in each scenario.

**Table S11. Runtime of each method calculated on 1,523 individuals and 17,619 genes of the large size European sample under the null hypothesis**. Note that if the analyses are conducted several times, with for instance different MAF thresholds or modes of inheritance, the pre-treatment part does not have to be performed again.

**Figure S1. Histogram of adjusted powers of the correction methods for the large size European sample (n=1,523) at the level *α*** = **0.001**. (A) Without stratification. (B) With moderate stratification. (C) With high stratification. Relative risks considered vary from 2 to 4 on the x-axis.

**Figure S2. Histogram of adjusted powers for the correction methods for the large size Worldwide sample at the level *α*** = **0.001**. (A) Without stratification. (B) With moderate stratification. (C) With high stratification. Relative risks considered vary from 2 to 4 on the x-axis.

**Figure S3. Histogram of powers for methods with a correct type I error rate for the large size European sample (n=1,523) at the level *α*** = **0.001**. (A) Principal components. (B) Linear Mixed Models. (C) LocPerm. Relative risks vary from 2 to 4 on the x-axis.

**Figure S4. Histogram of powers for methods with a correct type I error rate for the large size Worldwide sample at the level *α*** = **0.001**. (A) Principal components. (B) Linear Mixed Models. (C) LocPerm. Relative risks vary from 2 to 4 on the x-axis.

**Figure S5. Histogram of adjusted powers of the correction methods the small size sample at the level *α*** = **0.001**. (A) Scenarios with 50 cases from Southern-Europe. (B)Scenarios with 50 cases from the whole Europe. (C) Scenarios with 50 cases from the Worldwide sample. The relative risk is fixed at 4.

## Bibliography

1. Auer PL, Lettre G. Rare variant association studies: considerations, challenges and opportunities. Genome Med. 2015;7(1):16. doi: 10.1186/s13073-015-0138-2. PubMed PMID: 25709717; PubMed Central PMCID: PMCPMC4337325

2. Lee S, Abecasis GR, Boehnke M, Lin X. Rare-variant association analysis: study designs and statistical tests. Am J Hum Genet. 2014;95(1):5–23. doi: 10.1016/j.ajhg.2014.06.009. PubMed PMID: 24995866; PubMed Central PMCID: PMCPMC4085641

3. Morgenthaler S, Thilly WG. A strategy to discover genes that carry multi-allelic or mono-allelic risk for common diseases: a cohort allelic sums test (CAST). Mutat Res. 2007;615(1-2):28–56. doi: 10.1016/j.mrfmmm.2006.09.003. PubMed PMID: 17101154

4. Wu MC, Lee S, Cai T, Li Y, Boehnke M, Lin X. Rare-variant association testing for sequencing data with the sequence kernel association test. Am J Hum Genet. 2011;89(1):82–93. doi: 10.1016/j.ajhg.2011.05.029. PubMed PMID: 21737059; PubMed Central PMCID: PMCPMC3135811

5. Patterson N, Price AL, Reich D. Population structure and eigenanalysis. PLoS Genet. 2006;2(12):e190. doi: 10.1371/journal.pgen.0020190. PubMed PMID: 17194218; PubMed Central PMCID: PMCPMC1713260

6. Price AL, Patterson NJ, Plenge RM, Weinblatt ME, Shadick NA, Reich D. Principal components analysis corrects for stratification in genome-wide association studies. Nat Genet. 2006;38(8):904–9. doi: 10.1038/ng1847. PubMed PMID: 16862161

7. Kang HM, Sul JH, Service SK, Zaitlen NA, Kong SY, Freimer NB, et al. Variance component model to account for sample structure in genome-wide association studies. Nat Genet. 2010;42(4):348–54. doi: 10.1038/ng.548. PubMed PMID: 20208533; PubMed Central PMCID: PMCPMC3092069

8. Lippert C, Listgarten J, Liu Y, Kadie CM, Davidson RI, Heckerman D. FaST linear mixed models for genome-wide association studies. Nat Methods. 2011;8(10):833–5. doi: 10.1038/nmeth.1681. PubMed PMID: 21892150

9. Listgarten J, Lippert C, Kadie CM, Davidson RI, Eskin E, Heckerman D. Improved linear mixed models for genome-wide association studies. Nat Methods. 2012;9(6):525–6. doi: 10.1038/nmeth.2037. PubMed PMID: 22669648; PubMed Central PMCID: PMCPMC3597090

10. Zhou X, Stephens M. Genome-wide efficient mixed-model analysis for association studies. Nat Genet. 2012;44(7):821–4. doi: 10.1038/ng.2310. PubMed PMID: 22706312; PubMed Central PMCID: PMCPMC3386377

11. Jiang Y, Epstein MP, Conneely KN. Assessing the impact of population stratification on association studies of rare variation. Hum Hered. 2013;76(1):28–35. doi: 10.1159/000353270. PubMed PMID: 23921847; PubMed Central PMCID: PMCPMC4406348

12. Mathieson I, McVean G. Differential confounding of rare and common variants in spatially structured populations. Nat Genet. 2012;44(3):243–6. doi: 10.1038/ng.1074. PubMed PMID: 22306651; PubMed Central PMCID: PMCPMC3303124

13. O’Connor TD, Kiezun A, Bamshad M, Rich SS, Smith JD, Turner E, et al. Fine-scale patterns of population stratification confound rare variant association tests. PLoS One. 2013;8(7):e65834. doi: 10.1371/journal.pone.0065834. PubMed PMID: 23861739; PubMed Central PMCID: PMCPMC3701690

14. Liu Q, Nicolae DL, Chen LS. Marbled inflation from population structure in gene-based association studies with rare variants. Genet Epidemiol. 2013;37(3):286–92. doi: 10.1002/gepi.21714. PubMed PMID: 23468125; PubMed Central PMCID: PMCPMC3716585

15. De la Cruz O, Raska P. Population structure at different minor allele frequency levels. BMC Proc. 2014;8(Suppl 1 Genetic Analysis Workshop 18Vanessa Olmo):S55. doi: 10.1186/1753-6561-8-S1-S55. PubMed PMID: 25519390; PubMed Central PMCID: PMCPMC4143691

16. Moore CB, Wallace JR, Wolfe DJ, Frase AT, Pendergrass SA, Weiss KM, et al. Low frequency variants, collapsed based on biological knowledge, uncover complexity of population stratification in 1000 genomes project data. PLoS Genet. 2013;9(12):e1003959. doi: 10.1371/journal.pgen.1003959. PubMed PMID: 24385916; PubMed Central PMCID: PMCPMC3873241

17. Zawistowski M, Reppell M, Wegmann D, St Jean PL, Ehm MG, Nelson MR, et al. Analysis of rare variant population structure in Europeans explains differential stratification of gene-based tests. Eur J Hum Genet. 2014;22(9):1137–44. doi: 10.1038/ejhg.2013.297. PubMed PMID: 24398795; PubMed Central PMCID: PMCPMC4135410

18. Babron MC, de Tayrac M, Rutledge DN, Zeggini E, Genin E. Rare and low frequency variant stratification in the UK population: description and impact on association tests. PLoS One. 2012;7(10):e46519. doi: 10.1371/journal.pone.0046519. PubMed PMID: 23071581; PubMed Central PMCID: PMCPMC3465327

19. Luo Y, Maity A, Wu MC, Smith C, Duan Q, Li Y, et al. On the substructure controls in rare variant analysis: Principal components or variance components? Genet Epidemiol. 2018;42(3):276–87. doi: 10.1002/gepi.22102. PubMed PMID: 29280188; PubMed Central PMCID: PMCPMC5851819

20. Persyn E, Redon R, Bellanger L, Dina C. The impact of a fine-scale population stratification on rare variant association test results. PLoS One. 2018;13(12):e0207677. doi: 10.1371/journal.pone.0207677. PubMed PMID: 30521541; PubMed Central PMCID: PMCPMC6283567

21. Wang C, Zhan X, Bragg-Gresham J, Kang HM, Stambolian D, Chew EY, et al. Ancestry estimation and control of population stratification for sequence-based association studies. Nat Genet. 2014;46(4):409–15. doi: 10.1038/ng.2924. PubMed PMID: 24633160; PubMed Central PMCID: PMCPMC4084909

22. Baye TM, He H, Ding L, Kurowski BG, Zhang X, Martin LJ. Population structure analysis using rare and common functional variants. BMC Proc. 2011;5 Suppl 9:S8. doi: 10.1186/1753-6561-5-S9-S8. PubMed PMID: 22373300; PubMed Central PMCID: PMCPMC3287920

23. Sha Q, Zhang K, Zhang S. A Nonparametric Regression Approach to Control for Population Stratification in Rare Variant Association Studies. Sci Rep. 2016;6:37444. doi: 10.1038/srep37444. PubMed PMID: 27857226; PubMed Central PMCID: PMCPMC5114546

24. Zhang Y, Guan W, Pan W. Adjustment for population stratification via principal components in association analysis of rare variants. Genet Epidemiol. 2013;37(1):99–109. doi: 10.1002/gepi.21691. PubMed PMID: 23065775; PubMed Central PMCID: PMCPMC4066816

25. Zhang Y, Shen X, Pan W. Adjusting for population stratification in a fine scale with principal components and sequencing data. Genet Epidemiol. 2013;37(8):787–801. doi: 10.1002/gepi.21764. PubMed PMID: 24123217; PubMed Central PMCID: PMCPMC3864649

26. Listgarten J, Lippert C, Heckerman D. FaST-LMM-Select for addressing confounding from spatial structure and rare variants. Nat Genet. 2013;45(5):470–1. doi: 10.1038/ng.2620. PubMed PMID: 23619783

27. Genomes Project C, Auton A, Brooks LD, Durbin RM, Garrison EP, Kang HM, et al. A global reference for human genetic variation. Nature. 2015;526(7571):68–74. doi: 10.1038/nature15393. PubMed PMID: 26432245; PubMed Central PMCID: PMCPMC4750478

28. Boisson-Dupuis S, Ramirez-Alejo N, Li Z, Patin E, Rao G, Kerner G, et al. Tuberculosis and impaired IL-23-dependent IFN-gamma immunity in humans homozygous for a common TYK2 missense variant. Sci Immunol. 2018;3(30). doi: 10.1126/sciimmunol.aau8714. PubMed PMID: 30578352; PubMed Central PMCID: PMCPMC6341984

29. Mullaert J, Bouaziz M, Seeleuthner Y, Bigio B, Casanova J-L, Alcais A, et al. Taking population stratification into account by local permutations in rare-variant association studies on small samples. bioRxiv. 2020:2020.01.29.924977. doi: 10.1101/2020.01.29.924977

30. Moutsianas L, Agarwala V, Fuchsberger C, Flannick J, Rivas MA, Gaulton KJ, et al. The power of gene-based rare variant methods to detect disease-associated variation and test hypotheses about complex disease. PLoS Genet. 2015;11(4):e1005165. doi: 10.1371/journal.pgen.1005165. PubMed PMID: 25906071; PubMed Central PMCID: PMCPMC4407972

31. Belkadi A, Bolze A, Itan Y, Cobat A, Vincent QB, Antipenko A, et al. Whole-genome sequencing is more powerful than whole-exome sequencing for detecting exome variants. Proc Natl Acad Sci U S A. 2015;112(17):5473–8. doi: 10.1073/pnas.1418631112. PubMed PMID: 25827230; PubMed Central PMCID: PMCPMC4418901

32. Anderson CA, Pettersson FH, Clarke GM, Cardon LR, Morris AP, Zondervan KT. Data quality control in genetic case-control association studies. Nat Protoc. 2010;5(9):1564–73. doi: 10.1038/nprot.2010.116. PubMed PMID: 21085122; PubMed Central PMCID: PMCPMC3025522

33. Manichaikul A, Mychaleckyj JC, Rich SS, Daly K, Sale M, Chen WM. Robust relationship inference in genome-wide association studies. Bioinformatics. 2010;26(22):2867–73. doi: 10.1093/bioinformatics/btq559. PubMed PMID: 20926424; PubMed Central PMCID: PMCPMC3025716

34. Consortium UK, Walter K, Min JL, Huang J, Crooks L, Memari Y, et al. The UK10K project identifies rare variants in health and disease. Nature. 2015;526(7571):82–90. doi: 10.1038/nature14962. PubMed PMID: 26367797; PubMed Central PMCID: PMCPMC4773891

35. Sudlow C, Gallacher J, Allen N, Beral V, Burton P, Danesh J, et al. UK biobank: an open access resource for identifying the causes of a wide range of complex diseases of middle and old age. PLoS Med. 2015;12(3):e1001779. doi: 10.1371/journal.pmed.1001779. PubMed PMID: 25826379; PubMed Central PMCID: PMCPMC4380465

36. Rudolph PE. Good, Ph.: Permutation Tests. A Practical Guide to Resampling Methods for Testing Hypotheses. Springer Series in Statistics, Springer-Verlag, Berlin — Heidelberg — New York: 1994, x, 228 pp., DM 74,00; ōS 577.20; sFr 74.–. ISBN 3-540-94097-9. Biometrical Journal. 1995;37(2):150-. doi: 10.1002/bimj.4710370203

37. Zhou X, Stephens M. Efficient multivariate linear mixed model algorithms for genome-wide association studies. Nat Methods. 2014;11(4):407–9. doi: 10.1038/nmeth.2848. PubMed PMID: 24531419; PubMed Central PMCID: PMCPMC4211878

38. Zhang Y, Pan W. Principal component regression and linear mixed model in association analysis of structured samples: competitors or complements? Genet Epidemiol. 2015;39(3):149–55. doi: 10.1002/gepi.21879. PubMed PMID: 25536929; PubMed Central PMCID: PMCPMC4366301

39. Zhang X, Basile AO, Pendergrass SA, Ritchie MD. Real world scenarios in rare variant association analysis: the impact of imbalance and sample size on the power in silico. BMC Bioinformatics. 2019;20(1):46. doi: 10.1186/s12859-018-2591-6. PubMed PMID: 30669967; PubMed Central PMCID: PMCPMC6343276

40. Zhou W, Nielsen JB, Fritsche LG, Dey R, Gabrielsen ME, Wolford BN, et al. Efficiently controlling for case-control imbalance and sample relatedness in large-scale genetic association studies. Nat Genet. 2018;50(9):1335–41. doi: 10.1038/s41588-018-0184-y. PubMed PMID: 30104761; PubMed Central PMCID: PMCPMC6119127

